# Sparse coding models predict a spectral bias in the development of primary visual cortex (V1) receptive fields

**DOI:** 10.1101/2022.03.17.484705

**Authors:** Andrew Ligeralde, Michael R. DeWeese

**Affiliations:** Redwood Center for Theoretical Neuroscience, University of California, Berkeley, CA, USA; Biophysics Graduate Group, University of California, Berkeley, CA, USA; Department of Physics, University of California, Berkeley, CA, USA; Helen Wills Neuroscience Institute, University of California, Berkeley, CA, USA

**Author notes:** Current Address: Redwood Center for Theoretical Neuroscience, University of California, Berkeley, CA, USA.

## Abstract

It is well known that sparse coding models trained on natural images learn basis functions whose shapes resemble the receptive fields (RFs) of simple cells in the primary visual cortex (V1). However, few studies have considered how these basis functions develop during training. In particular, it is unclear whether certain types of basis functions emerge more quickly than others, or whether they develop simultaneously. In this work, we train an overcomplete sparse coding model (Sparsenet) on natural images and find that there is indeed order in the development of its basis functions, with basis functions tuned to lower spatial frequencies emerging earlier and higher spatial frequency basis functions emerging later. We observe the same trend in a biologically plausible sparse coding model (SAILnet) that uses leaky integrate-and-fire neurons and synaptically local learning rules, suggesting that this result is a general feature of sparse coding. Our results are consistent with recent experimental evidence that the distribution of optimal stimuli for driving neurons to fire shifts towards higher frequencies during normal development in mouse V1. Our analysis of sparse coding models during training yields an experimentally testable prediction for V1 development that this shift may be due in part to higher spatial frequency RFs emerging later, as opposed to a global shift towards higher frequencies across all RFs, which may also play a role. We also find that at least two explanations could account for the order of RF development: 1) high frequency RFs require more information to be specified accurately, and thus may require more visual experience in order to learn, and 2) early development of low frequency RFs improves the sparseness and fidelity of the visual representation more than early development of high frequency RFs.

**Author summary:** We are interested in how visual neurons learn representations of the natural world. In particular, we want to know whether certain visual features are learned by the visual cortex earlier in development than others. To address this question, we turn to a class of algorithms that can learn to represent natural scenes in a sparse fashion, with only a few neurons active at any given time (population sparseness). While sparse coding has been used extensively to model the response properties of neurons in the visual cortex, we use it here to arrive at a quantitative description of the way neurons might learn to encode visual information during development. We find that receptive fields (RFs) tuned to lower spatial frequencies develop earlier in our sparse coding models compared to high frequency RFs. If our prediction is accurate, such a description would provide a general framework for understanding the development of the functional properties of V1 neurons and serve as a guide for future experimental studies. It could also lead to new computational models that learn from input statistics, as well as advances in the design of devices that can augment or replace human vision.

## Introduction

A central goal of systems neuroscience is to establish a precise quantitative description of how neurons learn to encode sensory stimuli. Simple cells in the primary visual cortex (V1) have well-studied response properties [1–4] and therefore offer a useful model system for understanding how these representations of the visual world are learned during development. In this work, we use computational models of neural encoding to understand how V1 simple cells learn to represent the visual world from a stream of visual input. While many response properties of V1 simple cells can emerge before eye-opening without the need for visual experience (e.g., orientation selectivity and ocular dominance), observations of changes in receptive field (RF) properties that depend on the nature of the visual environment suggest that plasticity in V1 is experience-dependent [5]. Experimental evidence also shows that early postnatal visual experience is necessary for natural scene representation and discriminability in V1 [6].

The process of learning to encode visual information in V1 has been modeled as an unsupservised learning problem in which neurons adapt their tuning properties in order to optimize some objective function based on the statistical structure of stimuli in the natural environment. One coding principle that has proven to be useful for understanding sensory representations is sparseness, which posits that the neural population should not only maximize fidelity to input stimuli, but also minimize the number of active units (*L*_0_ population sparseness), or the amount of neural activity across the population (*L*_1_ population sparseness) [7]. Sparseness is an appealing concept for biological systems, both in terms of conserving metabolic costs and efficiently representing natural scenes, which have sparse structure [8]. Indeed, sparse coding models trained on natural image data to jointly optimize both fidelity to the input and sparseness have been shown to learn basis functions whose response properties replicate simple cell receptive fields (RFs) of V1 neurons [9–11].

Moreover, Hunt et al. have demonstrated that training sparse coding models with unnatural training images results in basis functions resembling the RFs that arise when animals are reared with abnormal visual input, suggesting that sparse coding is a feature of experience-dependent development [12]. Zylberberg and DeWeese show that sparse coding can account for the experimentally observed decrease in sparsity of V1 neural encodings over development in ferrets. They replicate this observation in SAILnet, a biologically plausible sparse coding model, by tracking the changes in sparseness of the learned representations throughout model training [13].

In this work, we analyze sparse coding models during training to answer the following question: do some types of basis functions develop sooner than others, and if so, which ones? Experimental work demonstrates that over the course of development, the distribution of frequency tuning of V1 neurons shifts towards higher spatial frequencies, and this shift requires visual experience [14, 15]. However, the question remains whether this shift is due to high spatial frequency RFs emerging later after the early development of low spatial frequency RFs, or whether there is a global shift during development across all receptive fields towards higher spatial frequencies. We find that the Sparsenet model [9] predicts the former to be true: low spatial frequency basis functions tend to emerge earlier in training, and high spatial frequency basis functions tend to emerge later. In fact, we observe the same behavior for the SAILnet model [11] of sparse coding, which implements leaky integrate-and-fire neurons and synaptically local learning rules, suggesting both that this result is a general feature of sparse coding and that it is biologically plausible.

## Methods

### Data and pre-processing

The sparse coding models used in this work are trained on 16×16 patches drawn from 35 images in the van Hateren database [16]. The original images are 1024 × rows 1536 columns, with pixel values linearly proportional to intensity. They are then pre-processed using the same procedure described in [17]. The full images are first transformed to log-intensity to account for background luminance [18]. The central 1024×1024 region of each image is extracted and the mean is subtracted to yield a pixel distribution that is roughly symmetric around zero. Each image is then whitened and lowpass filtered in the frequency domain by multiplying with the following filter:

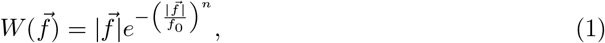

where 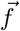 denotes the two-dimensional spatial frequency. The cutoff frequency *f*_0_ is set to 200 cycles/image, and the steepness parameter *n* is set to 4 to produce a sharp cutoff without introducing ringing in the space domain [19]. This filter is chosen to attenuate low frequencies while boosting high frequencies. For natural images, which typically have power spectra that roughly obey a 1/*f* ^2−*η*^ power law, with 0 < *η* < 0.3 [20], the filter flattens out the power spectrum to approximately whiten the images. Consistent with this, we find that the filter tends to overcompensate, giving slightly more power to higher frequencies than to lower frequencies (S1 Fig). The central 512 × 512 region of the image frequency domain is then extracted and inverse Fourier transformed to yield a 512×512 image that is down sampled by a factor of two from the original. The set of images obtained this way is then multiplied by a single scale factor so that the variance of the entire ensemble is 1.0. The filenames of the exact images used are listed in [17].

### Sparsenet simulation details

Sparsenet [9] is a sparse coding model that learns to approximate reconstructions of natural images as a linear combination of basis functions. The model is trained by alternately optimizing the basis functions and the coefficients of the linear combination [9, 21, 22]. The model used here consists of 2048 basis functions (about 10 × overcomplete relative to the effective dimensionality of the input image patches). The basis functions are initialized with Gaussian white noise. At each training iteration, a batch of 100 image patches is presented to the model. The model is trained for 10^5^ iterations with a sparseness parameter of λ = 1. All learning rates are held constant throughout training and are specified in the code used to generate these results, publicly distributed by Yubei Chen (https://github.com/yubeic/Sparse-Coding/).

### SAILnet simulation details

The SAILnet [11] model is a network of leaky integrate-and-fire neurons that learns approximate reconstructions of natural images based on neuronal spiking activity. The neurons receive excitatory feed-forward input from the image pixels weighted by their respective receptive fields (basis functions) and inhibit each other by recurrent connections. If the net input to a neuron exceeds its threshold value in response to a given image, it will emit some number of spikes; these spike counts are analogous to the coefficients in Sparsenet. The model is trained on Hebbian and anti-Hebbian rules similar to those used in [23], with the additional constraint that learning is localized to each synapse without information from any other synapses in the network. As with Sparsenet, the basis functions are initialized with Gaussian white noise, and the model is trained with batches of 100 images for 10^5^ iterations. To ensure stable convergence and sufficient diversity of the basis functions, the learning rates and model parameters must be appropriately tuned. Otherwise, the basis functions might only learn low-frequency features or they could exhibit what we will refer to as “fluidity” and continue to change indefinitely rather than converge to a final shape. The learning rate schedule is as follows: the initial learning rate in the original SAILnet code (linked below) for the first 10^3^ iterations; the initial learning rates reduced by a factor of 10 from that point until 5 × 10^4^ iterations; and tuned down by another factor of 10 from that point until the end of training. We set the sparseness parameter *p*, which modulates the target number of spikes per image, at *p* = 0.025, and *θ*_0_, the initial firing thresholds, at *θ*_0_ = 4.0. The SAILnet results are generated using code publicly distributed by Joel Zylberberg (http://www.jzlab.org/sailcodes.html).

### Classification of basis functions by frequency

Power spectra of the basis functions are computed by 2-dimensional discrete fast Fourier transform. We sample the power spectra for frequencies of up to 8 cycles per image (half the width and length of the basis functions) and bin by amplitude. Each basis function is assigned a class based on the maximum power frequency in its power spectrum: for each network, the lowest third of maximum frequencies is classified as low frequency, the middle third is classified as mid frequency, and the highest third is classified as high frequency.

## Results

### Sparsenet learns low frequency basis functions earlier

To study the rate at which basis functions are learned in Sparsenet, we train an overcomplete Sparsenet model on whitened natural image patches [9, 17]. We display the full learned Sparsenet dictionary in S2 Fig. For the model details and link to the code used, see Methods.

The basis functions in the model are initialized with Gaussian-distributed white noise. The extent to which a basis function is learned at a given time step in training *t* is measured by the degree of similarity to its final learned shape at the final training time step *T*. We quantify this using cosine similarity, the cosine of the angle between two vectors, between a given basis function at t and that same basis function at *T*. This is expressed as

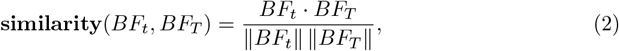

where *BF*_*t*_ denotes the basis function at a given time step *t, BF*_*T*_ denotes the final learned basis function, and ‖·‖ denotes the *L*_2_ norm. By definition, the maximum similarity between any two basis functions is 1, which indicates that they are equal up to a re-scaling of the pixel intensities. Two orthogonal basis functions have a similarity of 0.

We divide the basis functions into three equally sized categories — low, mid, or high frequency — based on the peak frequencies in their power spectra (see Methods for details). We plot the similarity over training time steps t and find that on average low frequency basis functions converge to their final learned shapes (reach similarity of 1) first, followed by the mid frequency basis functions, and the high frequency basis functions converging last (Fig 1A). The effect persists when binning the basis functions by the actual values of the peak frequency (number of cycles) in their power spectra, as opposed to binning into equally sized frequency batches, though differences in the number of basis functions for each category lead to higher variance for bins containing fewer basis functions (S3 Fig). These results demonstrate that the rate at which basis functions are learned is characterized by a spectral bias towards lower frequency features in the data.

**Fig 1.**
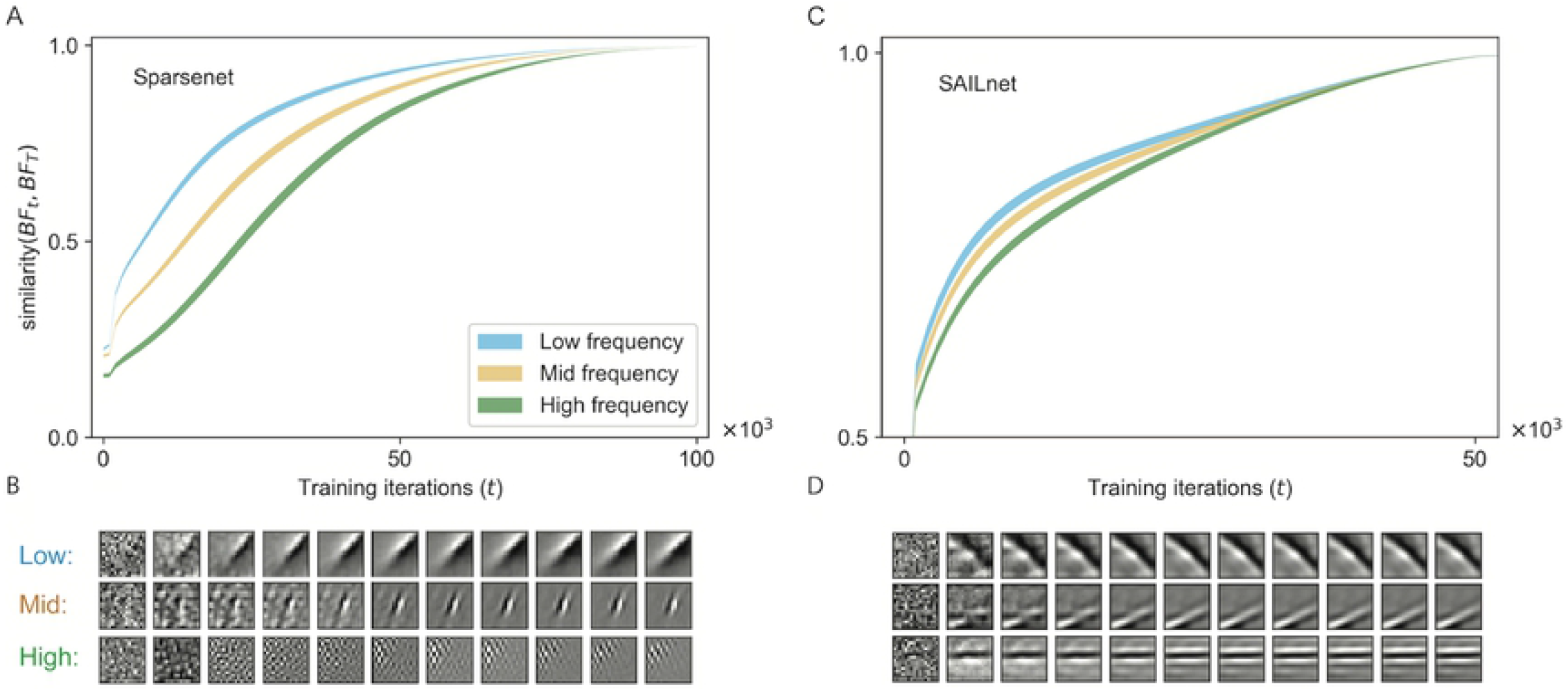
Low frequency basis functions develop earlier than high frequency basis functions in sparse coding models. (A) Convergence of basis functions over training time. A value of 1 on the y-axis denotes a basis function that has fully converged to its final learned shape *BF*_*T*_, so the higher the curve, the faster the basis function has converged. The shaded regions denote 95% confidence intervals about the mean for each category. (B) Representative examples of one basis function developing (left to right, shown every 1000 training iterations) from each frequency category, starting from random initialization. (C)—(D) Same analysis in SAILnet.

### A biologically plausible sparse coding model also learns low frequency basis functions earlier

We perform the same analysis as in the previous section using the SAILnet model, a biologically plausible sparse coding model that uses leaky integrate-and-fire (LIF) neurons and local learning rules [11]. Each neuron receives feed-forward input from image pixels, as well as inhibitory synaptic input from the other neurons in the network. The feed-forward weights and inhibitory synapses are both trained by iterative local learning rules — the update rule for an individual synaptic weight only depends on information available at that synapse during training, without requiring any information from any of the other synapses in the network. A third learning rule trains each neuron’s firing threshold, which modulates how often that neuron fires for a given amount of input. This rule is also local in that it only trains each threshold based on the current firing rate of that neuron, without access to the firing rates of any other neurons in the network. We emphasize these features of the model to demonstrate that this model achieves sparse representations while incorporating biologically plausible learning mechanisms — as could occur in real neuronal networks such as the population of simple cells in V1. For SAILnet, neural activities take on discrete values in terms of the number of spikes for a given input, and learning is local to synapses and neurons, neither of which is true for Sparsenet. For the model details and link to the code used, see Methods.

We train SAILnet on the same whitened natural image data as with Sparsenet for the same number of training iterations. We report the full learned SAILnet dictionary in S2 Fig. We find that on average, SAILnet also learns lower frequency basis functions earlier in training and higher frequency basis functions later in training (Fig 1C). Though the effect is smaller than in Sparsenet, the same order of development persists and also holds when binning by the actual peak frequency values (S3 Fig).

### Candidate explanations for fast learning of low frequency basis functions

There are several reasons one might expect low frequency basis functions to develop earlier than high frequency basis functions. We analyze two of these here and demonstrate that they are consistent with our findings.

### Higher frequency basis functions require more spatial precision to specify fully

The first potential reason for the observed spectral bias in training is that it requires more spatial precision to specify higher frequency basis functions, and therefore they may take more time to converge. We examine this by applying a small shift *ϵ* to the phase of each basis function along the direction of its maximum spatial gradient. We then compute **similarity**(*BF, BF* + *ϵ*), where *BF* + *ϵ* denotes the phase-shifted basis function, and find that high spatial frequency basis functions are changed more from their original shapes under *ϵ-*phase shifts (Fig 2A). The sensitivity of high frequency receptive fields to small perturbations suggests they may require more training data to converge to their final shapes.

**Fig 2.**
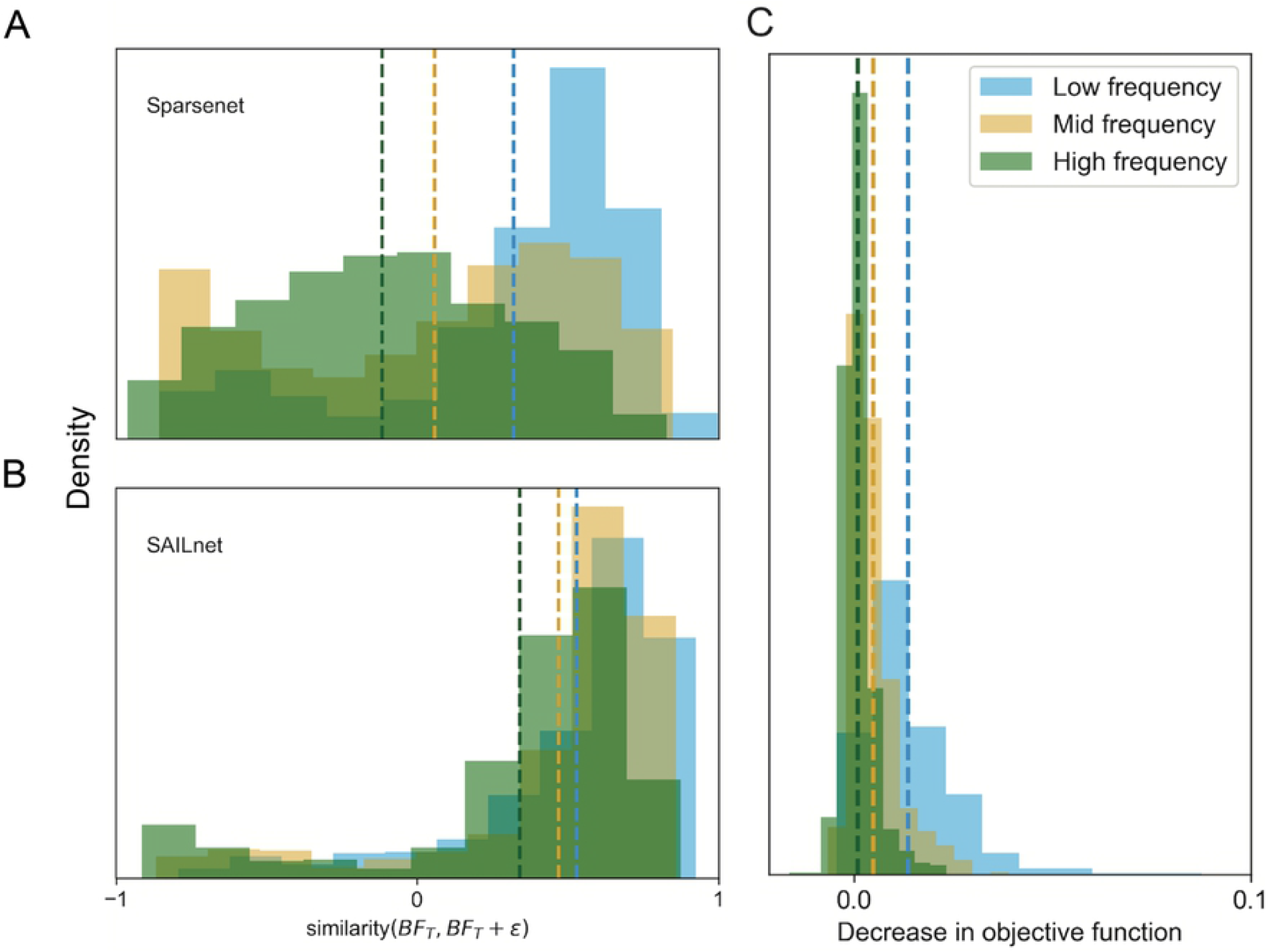
Two candidate explanations for early development of low frequency basis functions. (A)—(B) More information is needed to fully specify the shape of high frequency basis functions. To illustrate the sensitivity of a basis function’s spatial pattern to the parameters one might use to specify its detailed shape, such as its exact location, we computed the similarity between a given basis function and the same pattern shifted by a small amount. Phase shifts result in larger perturbations for higher frequency representations. The distribution of similarity between basis function and basis function under a 1-pixel phase shift for each frequency category. Vertical lines denote the means for each category. Note that for both models, the mean of the low frequency histogram is closer to 1 (exactly the same under phase shift) than that of the mid frequency or high frequency histograms, as one would expect. (C) Early development of lower frequency basis functions yields faster optimization of sparse coding objective function. For each of the three categories, the distribution of decreases in the Sparsenet objective function obtained by swapping one learned basis function from a fully converged dictionary into a randomly initialized model. Each objective function was evaluated on the same batch of 100 whitened natural image patches. A negative decrease corresponds to an increase in the objective function. Vertical lines denote the means for each category. Note that the low frequency histogram has a greater mean value than the mid frequency or high frequency histograms.

### Early learning of low frequency basis functions leads to faster optimization

Another reason we might expect low frequency basis functions to develop sooner is that they lead to faster optimization. We examine this possibility by comparing the model performance of a randomly initialized Sparsenet model to the performance of copies of the same model with one fully trained basis function swapped in for one of the randomly initialized basis functions. For each frequency category, we compute the distribution of changes in the objective function obtained by swapping in one fully trained basis function from that category. We find that swapping in lower frequency basis functions yields greater decreases in the objective than swapping in higher frequency basis functions (Fig 2B). This suggests that early development of low frequency basis functions leads to larger improvements in the sparse coding objective function than would early development of high frequency basis functions. Moreover, high frequency basis functions may only substantially improve sparse representations once low frequency basis functions have already emerged.

## Discussion

### The spectral bias in sparse coding model training is consistent with experimental findings in V1 development

In this work, we demonstrate that sparse coding models learn basis functions in a hierarchical manner: lower frequency basis functions are learned early in training, and higher frequency basis functions are learned later in training. Because Sparsenet and SAILnet represent two substantially different model architectures, we argue that our results are indicative of a general property of sparse coding, as opposed to being particular to a specific model architecture or optimization algorithm. In addition, because SAILnet is a biologically plausible model of sparse coding (see Methods), it further suggests that this observed spectral bias may be a feature of RF development in V1.

Our findings are consistent with experimental evidence that V1 becomes more attuned to higher spatial frequencies over the course of experience-dependent development. Chino et al. show a rapid increase in the mean optimal spatial frequency tuning of V1 neurons in macaque monkeys over the first four postnatal weeks [14]. Nishio et al. find that the overall distribution of optimal tuning of V1 neurons in mice shifts towards higher spatial frequencies from postnatal weeks 3-6. They also find that this shift in the distribution of optimal tuning towards higher frequencies does not occur for binocularly deprived animals, suggesting that visual experience is required for neurons to learn higher frequency representations [15].

Our results may provide additional insight into this phenomenon because we are able to observe the development of each individual basis function in the model, as opposed to just sampling from the distribution of tuning across the V1 neuronal population at different timepoints during training. In particular, we propose that this shift in the distribution is due to higher frequency receptive fields emerging later than the low frequency receptive fields. An alternative possibility is that it is due to a global shift across all receptive fields towards higher spatial frequencies during development. Future experimental work can help distinguish whether one or both of these explanations can account for the observations in [14, 15]. It may be experimentally challenging to track individual neuronal receptive fields over the full course of development, which would be the ideal way to distinguish whether high frequency RFs emerge late in development, as opposed to all RFs shifting to higher frequencies over time. Whether or not this can be done, it should be possible to sample from the population of receptive fields at various points in development and estimate the relative proportions of low, mid, and high frequency receptive fields at each time point. This could provide indirect evidence for one or the other of these possibilities, depending on the details of the distribution of RF shapes.

### Convergence of the SAILnet model depends strongly on initial conditions and model parameters

For some sets of hyperparameters and initial conditions, individual model neurons in SAILnet do not each converge to one final learned shape. Rather, individual neurons fluctuate, morphing from one shape to another throughout training; these fluctuations do not terminate even after training for many iterations. We refer to this phenomenon as fluidity. This is in contrast to Sparsenet, for which we have only observed smooth convergence of basis functions to their respective final shapes after many training iterations.

We note that when the basis functions are fluid during training, the metric of similarity over training time, and by extension, most simple metrics of convergence, are no longer meaningful, as there is no point at which every basis function has fully converged. Therefore, our main findings hold for a set of parameters and initial conditions that are sufficient to suppress fluidity (see Methods). Whether fluidity is a biological phenomenon is an interesting open question that we hope will be the subject of future experimental work.

### Basis function complexity and input data statistics could each account for spectral bias in RF development

We propose two candidate explanations for why lower frequency basis functions emerge earlier in sparse coding models. The first is a statement about complexity of the learned basis functions: higher frequency functions require more spatial precision to specify, and therefore may require more training data to accurately learn (Fig 1). The second is a statement about the dataset: early development of lower frequency basis functions leads to larger decreases in the sparse coding objective function, suggesting that the data contains more statistically relevant features at lower frequencies. As it is known that natural images exhibit a roughly 1/*f* ^2^ power spectrum [20], with more power concentrated at low frequencies, one might expect from the second mechanisms above that learning first lower frequencies on natural image data is better for optimization. However, we trained both models in this study on whitened natural images, which have approximately flat power spectra across all frequencies between our upper and lower frequency cutoffs. Moreover, our whitening procedure tends to overcompensate, boosting higher frequencies so power increases with frequency. This makes our results even more surprising: if anything, because of the dominance of higher frequencies in the dataset, we would expect the model’s spectral bias to lean towards learning higher frequencies first. This would seem to implicate the first mechanism over the second since the first mechanism is not so obviously dependent on stimulus properties.

It could also be the case that some aspects of higher-than-second-order statistics in the data not captured by the power spectrum may induce a bias towards learning lower frequencies early in training. Similar phenomena have also been observed in other statistical learning paradigms such as deep neural networks [24, 25], suggesting that spectral bias towards lower frequencies may be a general characteristic of representation learning.

Finally, there are many factors potentially affecting spectral bias in biological development that we do not consider in our modeling. For example, the optics of the eyes of young animals often change during development after eye opening so that spatial acuity increases over time [26]. Moreover, changes in the RFs of neurons upstream of V1, such as in the retinas or the lateral geniculate nucleus of the thalamus, are undoubtedly changing during development.

## Conclusion

We have demonstrated that sparse coding models learn low frequency basis functions earlier than high frequency basis functions during training and we propose this as an experimentally testable prediction for development of simple cell RFs in V1. Here, we only consider the development of V1 simple cells, rather than complex cells or neurons in higher visual areas. Other sparse coding models have the capacity to learn complex cell receptive field properties and topography [27]. We also do not consider excitatory connections between cells, which are a feature of the sparse coding model described in [28]. Future work could analyze development of the basis functions in these extensions of sparse coding. Other models of V1, such as the probabilistic Bayesian update model [29], would also be interesting to explore in the context of development.

Our spectral bias prediction was derived in the context of experience-dependent development, during which neuronal tuning adapts to natural scene statistics. It is possible that this particular order of development may not hold for experience-independent development, such as occurs in V1 prior to eye opening.This question could be addressed by considering different input data to the model. One possible input could be internally generated spontaneous neural activity, such as retinal waves, which play a role in the wiring of circuitry in early visual areas. For example, Dähne and colleagues implement slow feature analysis to encode retinal wave signals and find that the learned features correspond to the shapes of V1 complex cells [30]. Continuous-time implementations of sparse coding models trained on similar inputs could yield further insights into experience-independent development.

## Supporting information

**S1 Fig.**
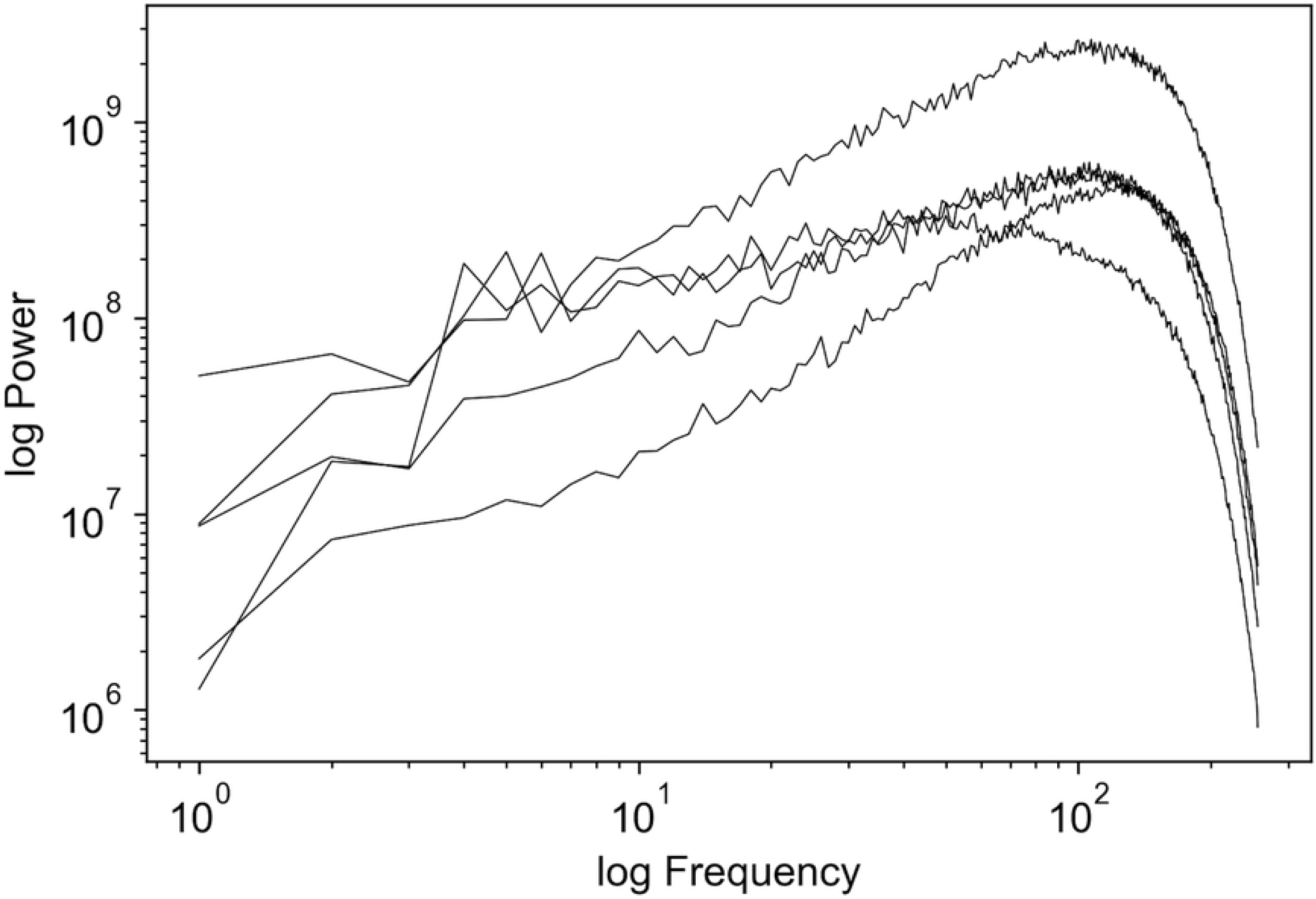
Power spectra of 5 randomly selected full, whitened images from dataset.

**S2 Fig.**
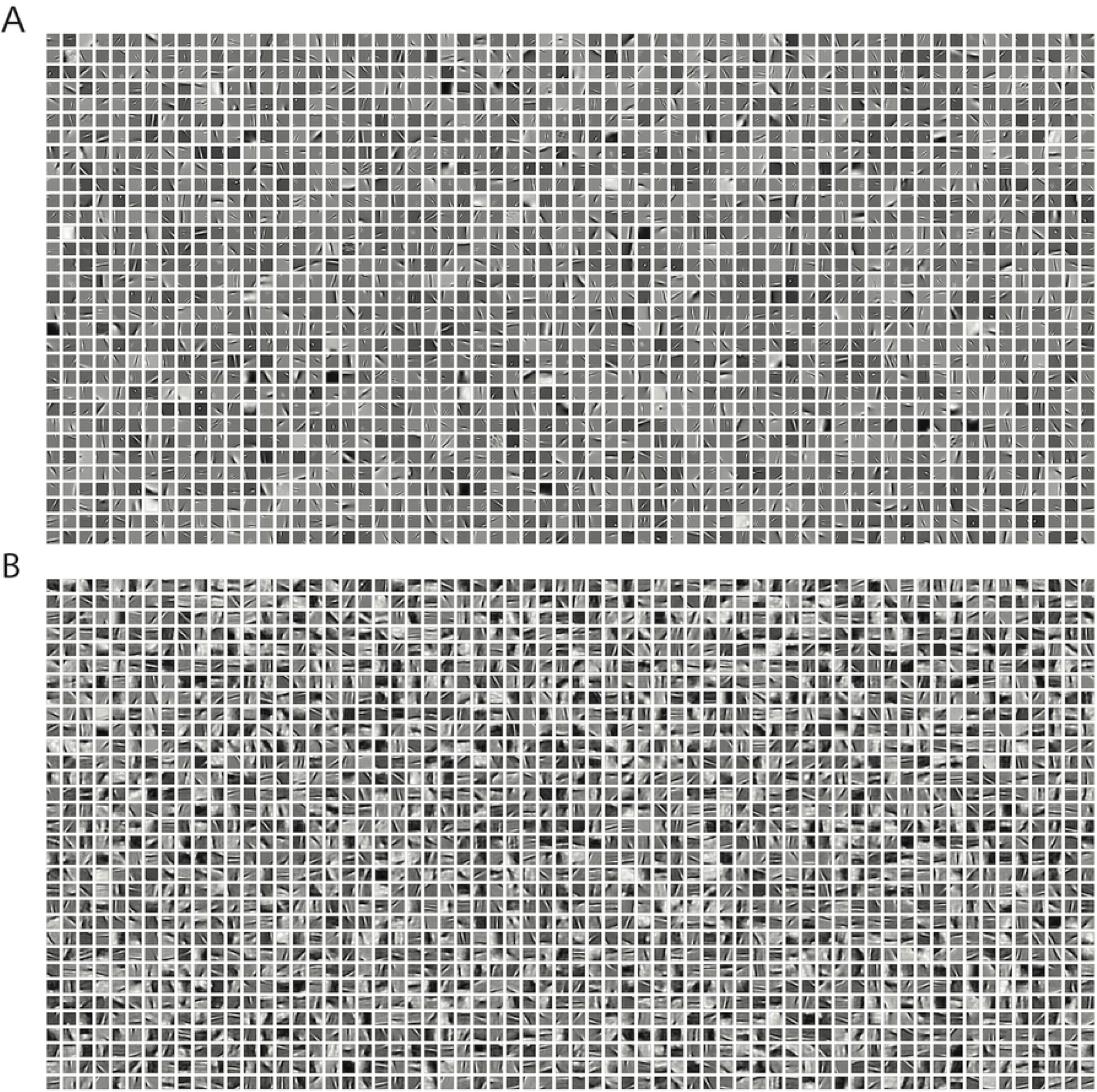
Complete dictionaries of learned basis functions. Both dictionaries are about 10x overcomplete (2048 basis functions) with respect to the dimensionality (200 features) of the in the image patches of the training set.

**S3 Fig.**
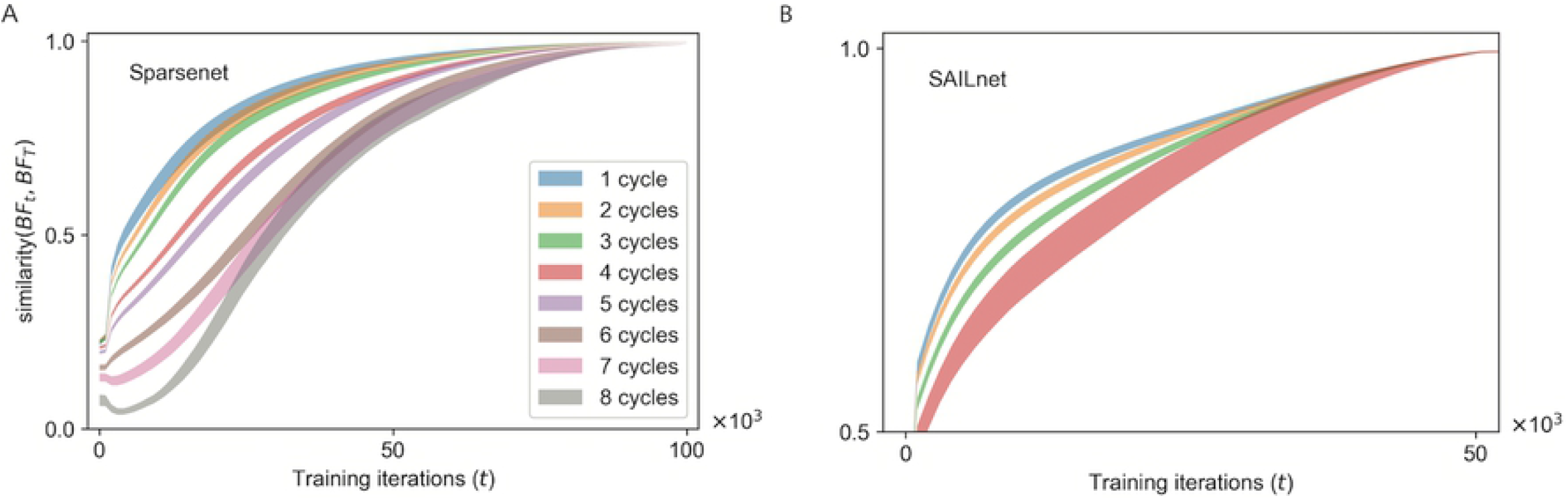
Direct correlation between number of cycles in peak frequency of basis function and time to develop.

## Acknowledgments

The authors thank Joel Zylberberg, Eric Dodds, Yubei Chen, and Bruno Olshausen for many helpful discussions.

